# Quality Assurance Strategies for Brain State Characterization by MEMRI

**DOI:** 10.64898/2026.04.10.717774

**Authors:** Taylor W. Uselman, Russell E. Jacobs, Elaine L. Bearer

**Affiliations:** University of New Mexico, School of Medicine, Albuquerque, NM; Zilkha Neurogenetic Institute, Keck School of Medicine, University of Southern California, Los Angeles, CA; Division of Biology and Biological Engineering, California Institute of Technology, Pasadena, CA

**Keywords:** Manganese-enhanced MRI (MEMRI), In vivo neuroimaging, Longitudinal imaging, Brain-wide mapping, Atlas segmentation, Statistical analysis, Simulation of noise-only images with and without investigator-embedded signals, Optimization of statistical mapping parameters to maximize accuracy

## Abstract

**Background:** Manganese-enhanced magnetic resonance imaging (MEMRI) is a powerful approach for mapping brain-wide neural activity and axonal projections *in vivo*. Yet standardized computational frameworks for voxel-wise and atlas-based characterization of brain states across large experimental cohorts remain limited.

**New method:** Here, we present methodological advances for preprocessing and statistical analysis of MEMRI datasets to support scalable, reproducible cohort-level analyses. Quality assurance metrics were developed to evaluate images, cohort-level anatomical alignment, and intensity normalization. Using simulated data, we optimized smoothing, effect-size, and cluster-size thresholds to balance sensitivity and specificity in voxel-wise statistical mapping. We developed ‘InVivoSegment’ software to apply to our new *InVivo* Atlas for segmentation of MEMRI data and interpretation of brain-wide activity.

**Results:** Quality assurance analyses established benchmarks for Mn(II)-induced signal- and contrast-to-noise evaluation, precise cohort-level alignment at 100 μm isotropic resolution, and robust intensity normalization. Balanced accuracy and Youden’s J statistics were calculated from simulated true positive and noise-only intensities, which defined optimal parameters for smoothing kernel, cluster-size and effect-size thresholds during voxel-wise mapping. Segmentation of simulated data demonstrated reliable transformation of voxel-wise results into regional summaries and identified secondary thresholds that minimize noise-driven artifacts.

**Comparison with existing methods:** Approach to optimize correction parameters for statistical mapping using simulated images improves voxel- and segment-wise sensitivity compared to FDR/FWE-based correction procedures.

**Conclusions:** These methodological advances enable scalable, reproducible, brain-wide quantification of longitudinal changes in MEMRI studies, strengthen mechanistic investigation of brain-state dynamics relevant to human health, and provide broadly applicable tools for neuroimaging analyses beyond MEMRI applications.

**Highlights:**

- Quantitative assurance of image quality complements visual assessment for cohort-level batch processing.
- Optimization of parameters using simulated noise-only images with and without investigator-embedded signal for voxel-wise mapping.
- A new software, “InVivoSegment” together with a labeled atlas, automates reliable user-friendly segmentation of voxel-wise data.
- Methodological advances in MEMRI data processing and computational analyses support scalable voxel- and segment-wise quantification of brain-wide neural activity.

## 1. Introduction

Computational analysis of images of the brain from MRI has transformed brain-wide neuroscience research. Standardization of preprocessing together with atlas-based analyses are fundamental for computational analysis. Automated accurate preprocessing facilitates subsequent reproducible mapping for comparisons of neural activity across anatomical regions, time, studies, species and modalities^1-12^. Innovation in neuroimaging methodology has occurred at a remarkable pace, requiring concurrent development of new specialized software and digital pipelines for data processing. One such methodology is manganese-enhanced MRI (MEMRI), which increases signal contrast, and enables longitudinal mapping of brain-wide neural activity and axonal projections^13-21^, alongside many other applications^22-27^. Typical MEMRI preprocessing procedure includes repurposing preexisting approaches used in other methods^14,15,28-30^, yet unique signal characteristics of Mn(II)-enhanced intensities introduce specific challenges for such repurposing and requires new approaches as well.

Among the various functional neuroimaging modalities, MEMRI offers a unique and powerful capability for mapping brain-wide neural activity longitudinally *in vivo*. After systemic delivery, Mn(II) distribution in the brain reaches a peak rapidly, 24-48h post-administration, followed by slow clearance, with a gradual return to baseline a week or so later^31,32^. Before and during this rise to that peak, activity-dependent uptake of Mn(II) ions into neurons provides a direct report of voltage-dependent calcium entry, and therefore neuronal activation, via hyperintense signals in T_1_-weighed MRI^33-36^. A significant advantage of MEMRI is the ability to highlight cumulative activity patterns over periods when animals are awake and freely moving. Hence, activity patterns correspond to ethologically relevant experiences outside the scanner^30,33,37^. With whole-brain coverage and high spatial resolution (e.g., 100 µm isotropic), MEMRI provides precise anatomical localization of distributed neural activity which enables voxel-wise anatomical registration of brain images across large cohorts over time. When applied to rodent models, brain-wide dynamics can be characterized across a series of experiences with experimental manipulation^31,38,39^, a feat often not possible in human studies.

Despite the inherent strengths of MEMRI, standardized processing-analysis pipelines comparable to those widely available for other approaches are underdeveloped. Mn(II)-enhanced signals require unique considerations for intensity normalization and anatomical alignment, which, if not carefully addressed, can introduce signal artifacts from non-activity-dependent sources^30,40,41^. Within preprocessing workflows quality assurance primarily involves visual inspection^42^, yet quantitative metrics can promote reliability, reproducibility and comparability across studies^11,43^. Moreover, studies often relied on *a priori* region-of-interest analyses or manually drawn regions for segmentation^29,44,45^ or were restricted to small cohorts, thus limiting comprehensive, brain-wide inference. Automated atlas-segmentation applied to brain-wide MEMRI data of multiple individuals can facilitate regional comparisons^31,38,46,47^. Results from voxel-wise mapping are susceptible to false positives and/or negatives due to preprocessing choices for smoothing kernels, statistical thresholds, and cluster-size^48-50^. We and others have worked towards addressing these issues by developing MEMRI-specific processing procedures and by validating analytical approaches for various biological applications^13,14,31,38,51,52^. Still, limited availability of tools and standardized procedures impede the broader application of MEMRI and preclude cross-study comparison.

The present study introduces methodological advances which enhance the scalability, reproducibility, and analytical depth of MEMRI data processing. We use our pre-existing datasets together with simulations to validate and expand our semi-automated computational pipeline for voxel- and segment-wise analysis of longitudinal MEMRI data^31,38,53,54^. We integrate quality assurance steps into our preprocessing workflow using both qualitative and quantitative metrics to minimize non-biological variability. We discuss inverse atlas-to-data registration to ensure accurate and consistent anatomical segmentation. We optimize voxel-wise statistical mapping procedures to balance detection sensitivity with biological specificity by accounting for impacts of smoothing, effect size and cluster size constraints. A key innovation is the development of new software tools to enable segmentation of MEMRI data via our *InVivo* Atlas from a high-resolution MEMRI image of a living mouse brain. These advancements enable robust and scalable assessment of MEMRI data both voxel- and segment-wise, thereby providing a framework for studies to track brain state dynamics and/or map neural projections across large-cohorts over time.

### 2. Methods

### 2.1. Sample Dataset and Code Availability

The data analyzed in this paper were obtained from were obtained by us^31,38^, and are posted in our publicly available longitudinal MEMRI dataset hosted on OpenNeuro^54^. In this report, analyses included only the pre- and post-Mn(II) images of standard, C57BL/6J mice (n = 11) sampled from 5 different weaning litters from JAX^55^. These images captured baseline, non-contrast-enhanced anatomy and Mn(II) accumulation patterns over a 23 h period, while mice were in their home cage. Note this timing captures a composite representation of neural activity integrated over the post-injection period rather than momentary activity during scanning. All MR images were acquired at Caltech on an 11.7 T vertical bore (89 mm) Bruker BioSpin Avance DRX500 small-animal MRI system (Bruker BioSpin Inc, Billerica, MA). For experimental details, please refer to our published reports^31,38,54^. All original animal procedures were conducted in accordance with NIH Guide for the Care and Use of Laboratory Animals^56^ and approved by the Institutional Animal Care and Use Committees (IACUC) at University of New Mexico and Caltech. No new animal experiments were performed. Sample size was determined via power analysis. Software code mentioned throughout this manuscript is maintained in publicly accessible GitHub repositories.

### 2.2. MR Processing Pipeline and Image Quality Assurance Assessments

Here, we present our newly enhanced semi-automated processing and analysis pipeline based on previously reported data. We redesigned our pipeline explicitly to perform quality assurance checkpoints, quantitative tools to ensure accuracy and precision of processing outcomes, and a new Python package for fast and reproducible segmentation of brain images using the *InVivo* Atlas – a high-resolution, Mn(II)-enhanced MR image of a mouse brain with anatomical delineation of 116 brain regions. A detailed diagrammatic overview of this pipeline is shown in **Fig. 1**.

**Fig. 1.**
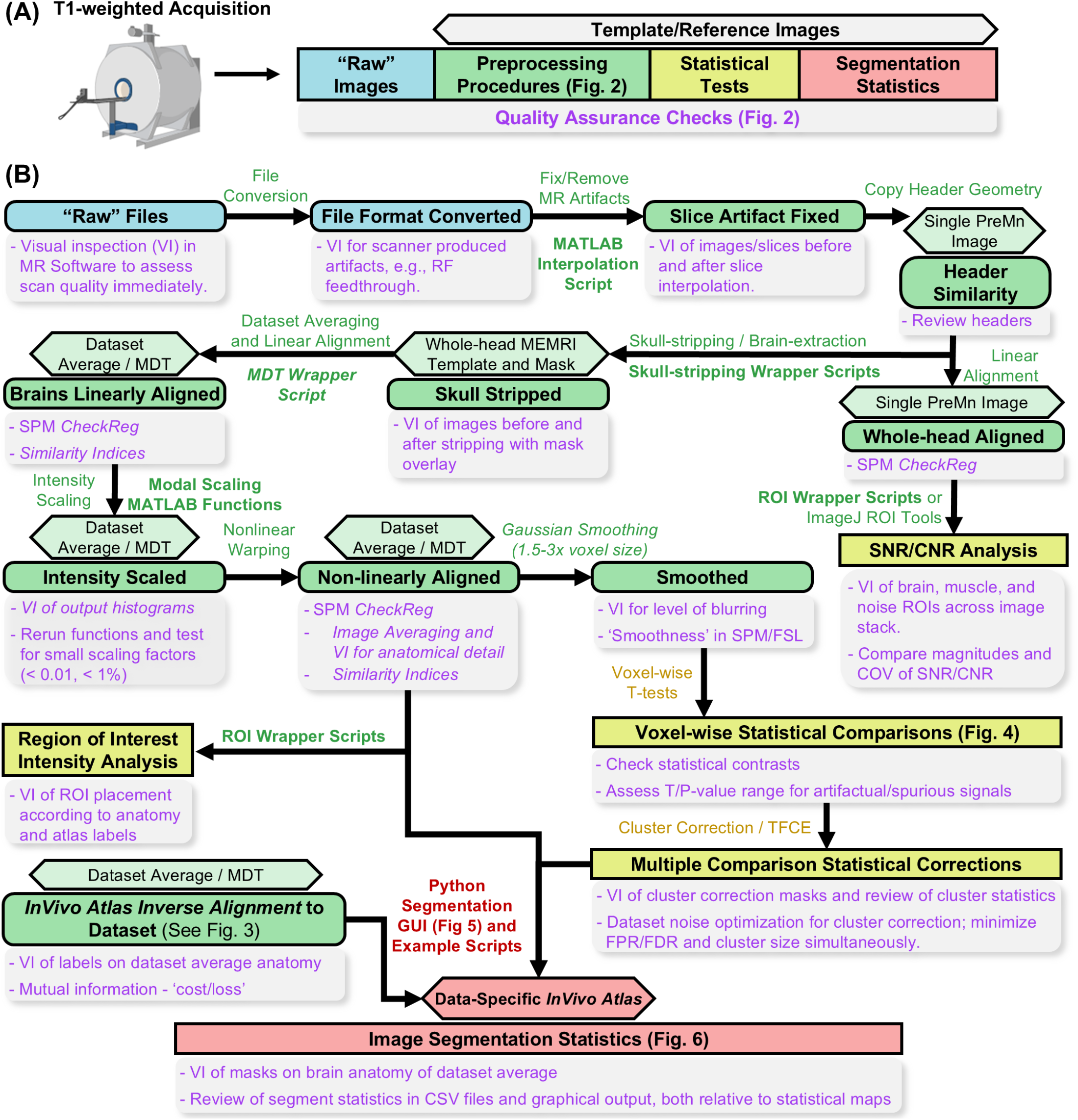
Diagram of the pipeline for MEMRI image processing and quality assurance at voxel- and segment-wise resolution. **(A)** A high-level overview of our acquisition, processing, and statistical analysis pipeline for MEMRI datasets. Each step of the process is color-coded in the diagram to identify the category listed in (B). Hexagonal shapes indicate whether a reference/template image is used, and if so, which image. Gray dropdowns with violet lettering indicate quality assurance assessments performed. MR Scanner was created with BioRender.com. **(B)** Color coding of steps: After collection of T_1_-weighted images (n = 11 at 2 conditions), MEMRI data stacks are processed in batch from ‘raw’ whole-head images (*top left*, blue) via typical image processing procedures (*middle*, green) for downstream analyses. Analyses include: 1) Statistical estimation and inference on signal intensity metrics (yellow); and 2) Anatomical segmentation and summarization of estimated statistics (*bottom*, red). Black arrows indicate flow of pipeline steps. Bolded labels indicate custom written code that facilitates automation.

#### 2.2.1. Raw image preprocessing and initial quality checks

After acquisition, images were visually inspected for large scale signal artifacts, such as RF feedthrough, aliasing, and other distortions using the scanner’s built-in viewing software (e.g., Bruker ParaVision). If identified immediately, images were re-acquired if longitudinal timing allowed. We converted raw Bruker images (2dseq) to NIfTI-1 format using ImageJ’s NIfTI plugin. After conversion, images were reviewed again for potential artifacts. Smaller RF-feedthrough slice artifacts were identified in a small number of images and removed via linear interpolation of adjacent slices using our custom MATLAB code^38,53^. After slices were fixed, header information was reviewed and the geometry (orientation, origin, voxel dimensions, units, s- and q-form matrices) of a single image was copied to all other images to ensure headers were identical^7^.

#### 2.2.2. Signal- and Contrast-to-Noise (SNR and CNR) measurements

We performed signal- and contrast-to-noise analyses to obtain a quantitative readout of overall image quality, consistency across the cohort, and to test for Mn(II) dosing. We perform a single rigid-body alignment with FSL flirt of all whole-head images to the single pre-Mn(II) image that was used as the header template to facilitate simultaneous measurement of the entire dataset. We measured signal intensities from within large areas of brain, muscle, and air for each image using the region-of-interest (ROI) manager in ImageJ. Average and standard deviations of signal were determined for each region in each pre- and post-Mn(II) image. We calculated image specific SNR and CNR as the average signal 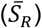 and standard deviation of signal (*σ*_*S*,*R*_) within a region (*R*) divided by the standard deviation of signal within non-tissue regions (*σ*_*S*,*Air*_), respectively 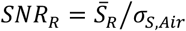 and *CNR*_*R*_ =*σ*_*S*,*R*_/*σ*_*S*,*Air*_). We also calculated Δ*SNR*_*R*_ and Δ*CNR*_*R*_ as the difference in SNR and CNR between pre- and post-Mn(II) images. The sample-wise coefficient of variation (COV) of SNR or CNR was defined as the standard deviation of SNR or CNR among either pre- or post-Mn(II) images divided by their respective average SNR or CNR. COV is a measure of variability in relation to the sample mean and is often used as a metric of precision for quality assurance purposes^57^.

#### 2.2.3. Monitoring skull-stripping, registration, and intensity normalization

Brain extraction was performed using our semi-automated skull-stripping procedure for mouse MEMRI data^13,38^. Remaining non-brain tissue was removed manually using the FSLeyes ‘edit’ tool. Skull-stripped brain images were anatomically aligned and intensity normalized to the dataset minimal deformation target (MDT), which was created from the dataset’s pre-Mn(II) images in an iterative align and average process^14^. We developed a new python wrapper script, which utilizes FSL functions, to generate the dataset MDT. We assessed MDT quality by visual inspection of anatomical detail in the average image at each step and quantitatively using Jaccard similarity indices (JSI) and normalized mutual information (NMI) comparing input images to the average at each step. JSI was calculated manually using *numpy* and Boolean logic, whereas NMI was calculated by applying *scikit-learn*’s built in NMI function on the numpy-generated brain intensity histograms each with 1000 bins and only for voxels above a minimum intensity threshold of 1000. The COV of these similarity indices were measured at each step to test for the convergence of brain anatomies from all individuals into the MDT.

After MDT creation, all pre- and post-Mn(II) images were rigid body aligned to the MDT using FSL *flirt*, and then intensity normalized via modal scaling^14,15,38,58^. Accurate and precise intensity normalization was assessed via visual inspection of grayscale intensity histograms and the COV of scaling factor magnitudes before and after scaling. Next, images were warped to the MDT using SPM12’s ‘Old Normalize’ function, which performs sequential affine and nonlinear transformations. JSI and NMI, and the COV of each, were calculated for each image before and after warping to quantitate anatomical similarity and hence alignment quality. Deformation vector fields from the warping were generated using the SPM12 ‘Deformations’ function on the warp field files generated from the SPM Normalize process. Using custom MATLAB scripts, we calculated the voxel-wise standard deviation of the deformation magnitude (SDDM)^59^, to asses voxel-wise anatomical variability across the dataset.

### 2.3. Voxel-wise Statistical Mapping of MR Data and Interpretation of Maps

Here, we performed statistical parametric mapping (SPM) paired t-tests in SPM12 comparing post-Mn(II) signals that are greater than pre-Mn(II) to detect Mn(II) enhancement. Images with known distributions of true and false positives signals were simulated to optimize required parameters for SPM of real MEMRI data, including: smoothing kernel full-width half maximum (FWHM)^5,60^, voxel-wise effect size and cluster size thresholds^48,49,61^.

#### 2.3.1 Simulation of false- and true-positive image matrices

Twenty-two 3D matrices (124 × 200 × 84 voxels; ML, AP, DV) were generated in R^62^ to match the dimensionality of our MEMRI images. Matrices were arbitrarily paired into two groups of n = 11, and exported to NIfTI^63^ ([1, 12], [2, 13], …, [11, 22]) (**Supplemental Fig. S1**). Voxel-wise noise was sampled independently from a Gaussian distribution derived from MEMRI SNR measurements (mean = 1000, sd = 300) (**Supplemental Fig. S1B-C**). We duplicated these 22 matrices to create a second set (23 - 44) for studies of true positive signals. A lattice (5 × 8 × 3) of voxel clusters, distributed ML/AP/DV, was defined within the matrices (**Supplemental Fig. S1D**). The size of voxel clusters within the lattice varied from 1 × 1 × 1 to 5 × 5 × 5 voxel cubes along the ML axis, and constant along the other two axes. Effect-sizes were similar within a cluster, varied from ***d*** = 0.546 to 0.806 to 0.950 between clusters along the DV axis, and were constant along the other two axes. This cluster-by-effect size pattern was repeated 8 times along the AP axis. Effect-sizes were chosen to correspond either to T_critical_ for a paired t-test at p < 0.05 and n = 11, or to the noncentrality parameters (NCP) for statistical power of 80 or 90%. To generate positive signals with these targeted effect sizes, we sampled intensities from a Gaussian distribution parameterized by the targeted mean difference ***µ***_Δ_ and voxel-wise signal variance ***v*** = (***σ***^2^ + ***ε***^2^) (**Supplemental Methods**). We simulated ***N*** = 10,000 random samples of size ***n*** = 11 from this distribution, calculated the effect size for each random sample set, and selected the sample of 11 that minimized the difference between simulated and targeted effect-size (max difference < 0.001) (**Supplemental Fig. S1E**). For visualization of the lattice within the matrix, we added a value of 6,000 arbitrary intensity units to the noise in each voxel cluster of the lattice in all 22 duplicated matrices (23 - 44). For half of the matrices (34 - 44), intensities from the selected sample set were added to the voxel clusters for each targeted effect-size (**Supplemental Fig. S1F**). Comparison of matrices represented true positives.

#### 2.3.2. Smoothing kernel, and effect- and cluster-size thresholds to optimize balanced accuracy of SPM results

Simulated image matrices, both noise-only and positives with noise, were smoothed with different sized Gaussian kernels: 0 μm, 150 μm or 300 μm FWHM. Following smoothing, SPM paired t-tests were performed separately on the n = 11 pairs in each simulated dataset: noise-only (1 - 11) versus noise-only (12 - 22); or noise-only plus lattice (23 - 33) versus positives (34 - 44). Resultant statistical maps were masked using pairwise combinations of cluster- and effect-size thresholds, set to match those of the embedded positive signals. For the comparison between noise-only matrices, the resultant statistical map displayed only false-positives; and for the comparison with embedded positive signal, the statistical map should give true positives within the lattice and false positives elsewhere. For the second comparison, we counted the number of ‘actual’ embedded positives, the number of ‘SPM-predicted’ positives (suprathreshold voxels), and the total number of voxels using an FSL wrapper script. From these voxel counts we computed the full confusion matrix, including false positive rate (FPR), false negative rate (FNR), specificity, sensitivity, balanced accuracy and Youden’s J^64,65^. Results were graphed in R using ggplot2^66^. We determined the optimal parameter set of smoothing kernel size, and effect- and cluster-size thresholds to be that which met SPM smoothness assumptions^5^, maximized balanced accuracy, and resulted in the greatest ratio of false positives removed over false negatives added.

#### 2.3.3. Voxel-wise variability and standardized effect-size maps

For voxel-wise mapping of Mn(II) accumulation, we performed paired SPM t-test in SPM12^60^ using the set of optimized parameters determined from simulations above; FWHM = 150 μm; d ≥ 0.546 (T_(10)_ = 1.812, p < 0.05); and cluster size ≥ 8 voxels. We calculated a “standard deviation” image by taking the square root of the residual square error output from SPM. We calculated a Cohen’s D map by dividing the paired T-statistic image by the square root of the sample size 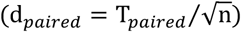. Statistical significance maps were generated by applying effect- and cluster-sizes thresholds. We mask all statistical maps to exclude non-brain voxels using a binarized version of the MDT.

### *2*.*4 InVivo* Atlas Segmentation

The *InVivo* Atlas is a high resolution (80 μm isotropic) grayscale MEMRI image^67^ with a corresponding label image for 116 brain regions. Segments were based on a combination of the Allen Institute’s CCF3 and Paxinos-Franklin histologically derived anatomy and naming^12,68^, and anatomical contrast in the MR image. Here, we developed the InVivoSegment software, which uses the *InVivo* Atlas aligned to a dataset to calculate segment-wise summaries of voxel-wise intensities. Here, we apply this software to three datasets: 1) a simulated image to validate segment measure calculations (from **2.4.2**. below); 2) statistical maps of simulated false and true positive comparisons (as in **2.3.1**. and **2.3.2**. above); 3) a Cohen’s d effect size map of real MEMRI data (as in **2.3.3**. above).

#### 2.4.1. Inverse alignment of *InVivo* Atlas to dataset

Inverse alignment of the high resolution *InVivo* Atlas to our dataset MDT was implemented in a multistep procedure: (1) harmonization of header geometry between the atlas images and the MDT; (2) stepwise linear and nonlinear forward alignment of the MDT to the atlas grayscale image; (3) inversion of the resulting transformation matrices and deformation fields; (4) application of the inverted transformations to the atlas grayscale image using trilinear or B-spline interpolation; and (5) application of the inverted transformations to the atlas label image using nearest-neighbor interpolation to preserve discrete segment boundaries. This procedure was implemented using multiple standard neuroimaging registration frameworks, including FSL, SPM, and ANTs^7,9,60^. Optimized registration parameters and scripts are provided for SPM *Normalize*, FSL *FNIRT*, and ANTs implementations. Alignment accuracy was assessed qualitatively by visual inspection and quantitatively by the magnitude and variance of JSI and NMI from comparisons of the MDT and warped images to the aligned grayscale atlas, respectively. Plots were generated in Excel.

#### 2.4.2. Creating a simulated image for segmentation validation

To validate segmentation accuracy, we created a single blank image with the same dimensionality and resolution as the MDT. We inserted four 7 × 7 × 7 voxel cubes of signal into three different anatomical segments: 1 in ACA; 1 in CP; and 2 in PRN. Signals within cubes followed a gradient pattern, with a minimum intensity of 1 in the outermost layer of the cube and a maximum intensity of 4 in the center voxel. We calculated the “ground truth” segmental measures for this simulation image in Excel, which was used for comparison to results from InVivoSegment processing (as in 2.4.3. below).

#### 2.4.3. Segmental measures of voxel-wise intensities extracted by the InVivoSegment software

Application of the segmented *InVivo* Atlas labels to simulated and real MEMRI data was performed using the “InVivoSegment” Python software package. For each atlas-defined segment, voxel-wise intensities intersecting the aligned segment label were extracted and summarized into segment-level metrics (segmental measures), which were exported in comma-separated value (CSV) format for downstream statistical analysis. To facilitate user accessibility, “InVivoSegment” includes a graphical user interface (GUI). Segmental measures of segmental intensities and volumes offered in this software include Mean, Median, standard deviation (StDev), first and third quartiles (Q1 and Q3), minimum (Min), maximum (Max), activation volume (number of suprathreshold voxels, ActVol), fractional activation volume (fraction of suprathreshold voxels relative to total segment volume, FAV), and the unweighted and signal-weighted 3D centroids. These metrics were calculated from both statistical maps from comparison of noise-only versus noise-only simulated data and from real MEMRI data. For each InVivoSegment run, CSV outputs were imported and indexed by atlas segment ID, anatomical domain, group, condition, image/subject identifier, and threshold value (if any). Additional measures (COV of effect size, and Euclidean Distance of centroid displacement) were calculated from the original 10 using Python in Jupyter notebooks. The segmental FAVs from InVivo segmentation of the noise-only statistical map were fitted to a Beta-distribution and the 99^th^ percentile was applied as a “noise-independent” secondary threshold for FAV in real MEMRI data. All other metrics were set to 0 for segments with FAV below this threshold, and then plotted as column graphs using InVivoSegment utilities^69,70^. Segment ordering followed the hierarchical domain structure provided by the *InVivo* Atlas sorting table (**Supplemental Table S1**).

## 3. Results

### 3.1. Quality assurance for longitudinal MEMRI datasets

To ensure image quality, we recommend all MEMRI images undergo multiple quality assurance (QA) steps throughout (**Fig. 1**). Here, we show examples of our preprocessing and QA. Raw images were visually inspected for MR acquisition artifacts, such as radiofrequency (RF) feedthrough, aliasing, and distortions (**Fig. 2A**). To test for robust Mn(II)-enhancement and consistent Mn(II) dosing between individuals, we recommend comparing SNR and CNR. As an example, we measured intensities in brain and muscle tissue, and in non-tissue (empty space) (**Fig. 2B**). We compared between images with and without Mn(II). If Mn(II) enhancement were robust and dosing consistent, we would expect SNR and CNR to increase with Mn(II) and be similar across images. From pre- to post-Mn(II), SNR increased in both brain (+4.0, p < 0.011) and muscle (+3.1, p < 0.044) (**Fig. 2B2**). CNR increased primarily in brain (+1.5, p < 0.0001) with a marginal increase in muscle (+0.5, p < 0.069) (**Fig. 2B3**). We found COV for both SNR and CNR to be low (COV < 0.3), demonstrating consistent enhancement across the dataset (**Fig. 2B4**). These initial checks ensure image quality for subsequent processing.

**Fig. 2.**
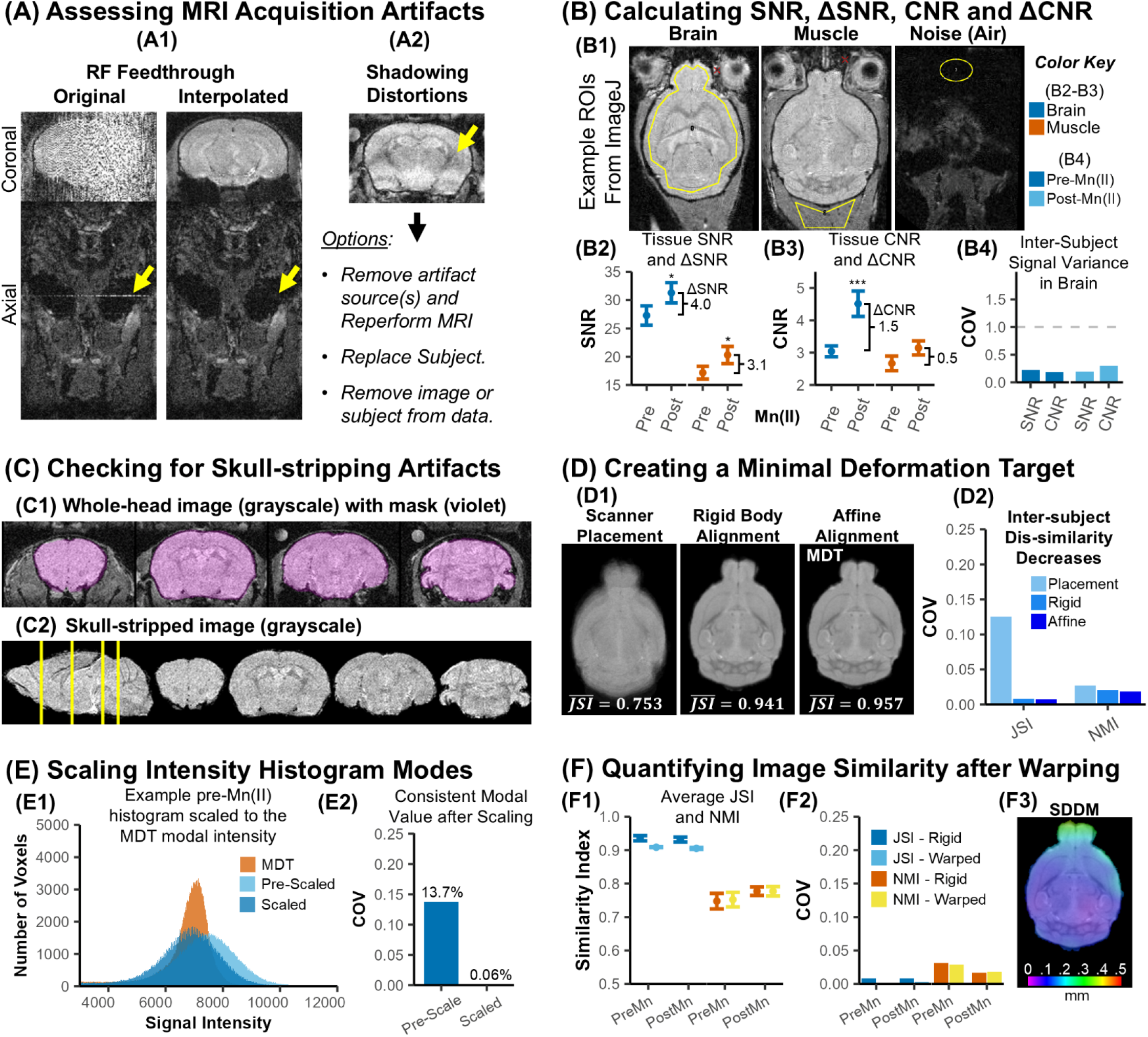
Quality assurance. **(A)** Visual inspection of raw data. Examples of two types of MR acquisition artifacts: **(A1)** radiofrequency (RF) feedthrough shown in coronal and axial slices (*left*, yellow arrow), which were removed via slice interpolation (*right*); and **(A2)** shadowing distortions. **(B)** Determining overall image quality by SNR and CNR. **(B1)** Regional intensity measurements in axial slices (yellow): Brain (*left*), Muscle (*middle*) and air (noise, *right*). Average **(B2)** SNR, **(B3)** CNR in brain (blue) and muscle (orange) and their change (ΔSNR/CNR) between pre- and post-Mn(II) images. **(B4)** COV of SNR and CNR in brain measurements for pre-Mn(II) (dark blue), and post-Mn(II) images (light blue). **(C)** Determining skull-stripping quality. Coronal slices from an example whole-head grayscale image **(C1)** with skull-stripping mask (violet) overlaid and **(C2)** after skull-stripping. **(D)** Measuring the convergence of data into the MDT. **(D1)** Axial slices of the average pre-Mn(II) image based on placement in the scanner (*left*), after rigid-body alignment (*middle*), and after an affine alignment (*right*). **(D2)** COV of similarity metrics at each step of MDT creation process. **(E)** Determining the accuracy and consistency of intensity normalization via modal scaling. **(E1)** Grayscale intensity histograms for the MDT (orange) and a single pre-Mn(II) image before (light blue) and after (dark blue) modal scaling. **(E2)** COV of scaling factor magnitudes. **(F)** Quantification of anatomical alignment for reliable voxel-wise comparisons. **(F1)** JSI (blues) and NMI (yellow/orange) of pre- and post-Mn(II) images after increasingly precise alignment. **(F2)** COV of similarity metrics after rigid alignment versus warping. **(F3)** Axial slice of the voxel-wise standard deviation of the deformation magnitude (SSDM) image; color coding in 0.1 mm (1 voxel) intervals. Error bars represent mean ± SEM. Paired t-tests, * p < 0.05, *** p < 0.001; n = 11.

A first step is to isolate the brain image by masking non-brain voxels^13^. Skull-stripped images resulting from our algorithm should be visually examined and manually masked of remaining non-brain voxels (**Fig. 2C**). These images are then aligned into a minimal deformation target (MDT) to be used as a template for further processing. To determine skull-stripping accuracy and MDT quality, we measured the Jaccard Similarity Index (JSI) and Normalized Mutual Information (NMI) between input images at each step in the MDT-generation process and the final MDT (**Fig. 2D**). We would expect JSI and NMI to increase in magnitude and decrease in variance. JSI increased from 0.753 from scanner placement alone to 0.957 after affine alignment, whereas NMI increased from 0.761 to 0.771. Sample-wise COV for JSI and NMI decreased to minimal levels after affine alignment (COV < 2% for each). Together these metrics reveal accurate isolation of brain anatomy, as well as strong anatomical convergence into a high-quality MDT.

A next step, following a rigid body alignment of skull-stripped images to the MDT, is to normalize grayscale intensities via modal scaling^14^. To test the accuracy and precision of intensity normalization, we inspected image intensity histograms for alignment of modal intensities to that of the MDT and by measuring the scaling factors magnitude and variability (**Fig. 2E**). Near-zero scaling factor magnitudes and low variability after scaling would indicate successful normalization. Histogram modes exhibited strong agreement with that of the MDT, as average scaling factor magnitudes decreased from 1.3% before scaling to 0.07% after, and COV of scaling factor magnitudes decreased from 13.7% before scaling to 0.06% after. These results demonstrate accurate and precise normalization of grayscale intensities across the dataset.

A final step is to warp individual brain anatomies into that of the MDT through successive affine and nonlinear alignments. To evaluate warping accuracy, we calculated JSI and NMI for each image and tested for decreased variability after warping (**Fig. 2F1**). Notably, JSI decreased in magnitude after warping (F_(1,40)_ = 127, p < 0.0001), while NMI magnitudes remained similar (F_(1,40)_ = 0.344, p < 0.561). This decrease in magnitude stemmed from signal artifacts around the brain surface after warping, which influenced the threshold-based masking for JSI calculations. Therefore, we advise against comparison of average similarity magnitudes after warping. Alternatively, sample variance for JSI and NMI magnitudes should decrease as anatomies converge. Indeed, COV for both metrics decreased to negligible levels after warping (COV_JSI_< 0.3%; COV_NMI_< 2.9%) (**Fig. 2F2**). Anatomical variability within a dataset may also influence alignment accuracy^59^. To determine the degree of anatomical variability and risk of misalignment, we calculated the voxel-wise standard deviation of deformation magnitude (SSDM) (**Fig. 2F3**). We found the brain-wide average SSDM to be ∼0.115 mm (just over one voxel). Thus, even a single rigid transform, enabled strong alignment. Together, these outcomes reveal that brain anatomies in our dataset are warped to the dataset MDT with high accuracy and are adequately prepared for subsequent statistical analyses.

### 3.2. Smoothing, effect- and cluster-sizes influences sensitivity and specificity of voxel-wise SPM

Statistical parametric mapping (SPM) has tradeoffs between specificity and sensitivity. First, spatial smoothing is a requirement of SPM^5,60^, yet excessive spatial smoothing reduces resolution and thereby spatial specificity. Second, insufficient correction for multiple voxel-wise comparisons leads to inflated false positive rates, yet typical correction procedures (e.g., FWE/FDR) may only detect signals with large effect sizes and/or cluster sizes. We systematically evaluated error rates after SPM paired t-tests on simulated data with and without positive signals embedded in discrete locations of a lattice of voxel clusters, using different combinations of smoothing kernel sizes (0, 150 and 300 μm), T-value thresholds (equivalent to the effect sizes, d = 0.546, 0.806, 0.950), and cluster sizes (1^3^ - 5^3^), the latter two corresponding to the simulated positive signals (**Fig. 3A**).

**Fig. 3.**
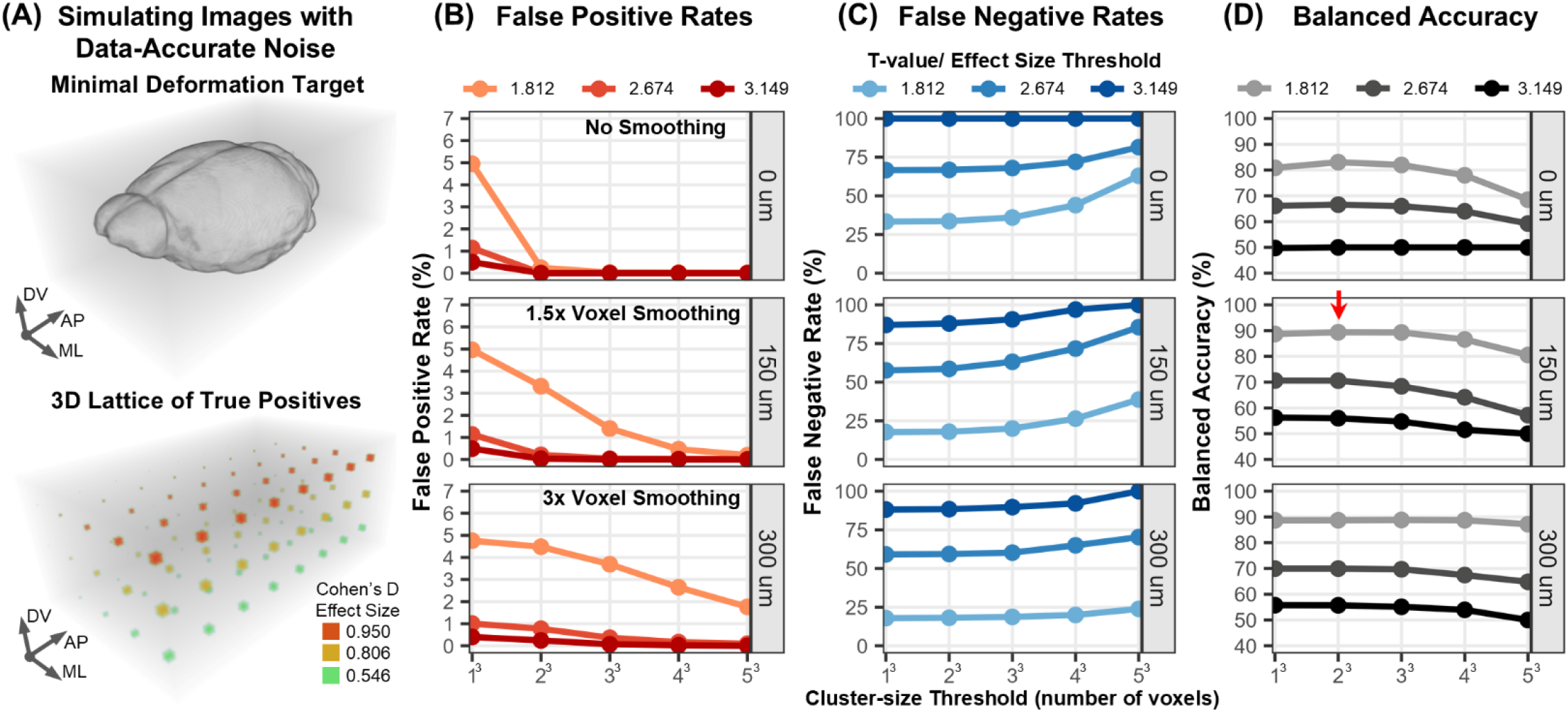
Considerations for smoothing, effect size and cluster size on sensitivity and specificity of statistical mapping. **(A)** Simulation of true positive signals at different effect- and cluster-sizes with data-accurate noise. Shown in perspective are 3D renderings of the dataset MDT (*top*) and true positive signals (*bottom*) in a 3D lattice of cubic voxel clusters and embedded in a matrix of noise (gray background), which matched the FOV/resolution of the whole-head MR data. In the lattice of true positives, cluster-size (number of voxels in a cube) varies along the dorsal-ventral axis (DV, 1^3^ - 5^3^voxels), effect-size varies along the medial-lateral axis (ML, d = 0.55 - 0.95), and this cluster-by-effect-size pattern was repeated 8 times along the anterior-posterior axis (AP). **(B-D)** Optimizing cluster and effect size-based thresholding to balance specificity and sensitivity of smoothed SPM results. Shown are line graphs of **(B)** False Positive Rates (FPR), **(C)** False Negative Rates (FNR), and **(D)** Balanced Accuracy for results of SPM paired t-tests on simulated data with different parameter combinations: No smoothing (*top*, 0 μm), 1.5x voxel smoothing (*middle*, 150 μm), and 3x voxel smoothing (*bottom*, 300 μm). T-value SPM thresholds correspond to effect sizes degrees of statistical power for a sample size of n = 11: 1) the minimum T_critical_ for p < 0.05 uncorrected (d = 0.546, T_(10)_= 1.812); 2) the noncentrality parameter (NCP) corresponding to 80% power at p < 0.05 (d = 0.806, T_(10)_= 2.674); and 3) NCP corresponding to 90% power at p < 0.05 (d = 0.950, _(10)_ = 3.149). NCP values were determined in G*Power 3.1^99^. The red arrow points to the parameter combination that maximizes balanced accuracy.

We examined consequences of the size of smoothing kernel on error rates in these statistical maps simulated to match the 100 μm isotropic voxels of our images. We aimed to minimize the rates of both false positives (FPR) outside and false negatives (FNR) inside the lattice (**Fig. 3A**, *bottom*) and maximize balanced accuracy, the average of specificity and sensitivity. The lattice allows us to test 3 different smoothing kernel sizes, 3 different T-values, and 5 different cluster-sizes. FPR was as expected (p < 0.05), for all combinations of kernel sizes and SPM thresholds for T-value and cluster-size (**Fig. 3B**). False negatives are when embedded positive signals are not detected by SPM. FNR was less than 20% for kernels of 150 μm and 300 μm but reached 33% when data was not smoothed (**Fig. 3C**). Further increasing smoothness (300 μm, FWHM) decreased spatial specificity without a meaningful improvement in detection. Thus, the optimal choice of smoothing appears to be the minimum required to meet SPM assumptions (150 μm, FWHM).

To test how well FPR corrections maintain sensitivity (1 - FNR), we measured error rates after setting different T-value or cluster-sizes on statistical maps of simulated data smoothed with a 150 μm kernel. Increasing T-value corresponding to d = 0.546 to 0.950 with cluster size = 1, decreased FPR 10-fold from 5% to 0.5% and increased FNR 4.8-fold from 18% to 87% (**Fig. 3B-C**). This represents a 1.6-fold decrease in balanced accuracy from 89% to 56% (**Fig. 3D** and **Supplemental Table S2**). Meanwhile, increasing cluster-size, from 1 to 125 voxels with effect-size constant at d = 0.546, decreased FPR 25-fold from 5% to 0.2% and increased FNR 2.2-fold from 18% to 39%. These thresholds applied to our simulated data resulted in a minimal 1.1-fold decrease in balanced accuracy, from 89% to 81%. Thus, cluster-based correction maintained sensitivity better than effect-size thresholding. A minimal T-value threshold of T = 1.812 together with a cluster-size of 8 was the single best choice for our data (**Fig. 3D**, red arrow), as it maximized the relative number of false positives removed (33%) over false negatives added (1.5%), and resulted in the greatest overall balanced accuracy in our data (89.4%).

Thus, our results suggest the following three steps: 1) apply the minimal 1.5x voxel smoothing kernel to meet SPM assumptions and to lessen the influence of voxel-wise noise; 2) use the minimal effect-size threshold to meet FPR and FNR used for an *a priori* power analysis; and 3) apply a cluster-size threshold that is close to the resolution of the data, e.g., 8 - 27 voxels for data smoothed by a 1.5x voxel smoothing kernel.

### 3.3. Statistical summaries for brain-wide, voxel-wise statistical mapping

There are several useful statistical outputs for characterizing voxel-wise intensities. These outputs include grayscale intensity images (**Fig. 4A**), averaged intensities of pre- and post-Mn(II) images (**Fig. 4B**), averaged voxel-wise differences in signal intensity (ΔSI) between pre- and post-Mn(II) images (**Fig. 4C1**), voxel-wise root-mean square error of ΔSI (**Fig. 4C2**), voxel-wise Cohen’s D effect sizes of ΔSI (**Fig. 4C3**), and statistical significance map of ΔSI between conditions (**Fig. 4C4**). This latter statistical significance map (**Fig. 4C4**) was generated by applying the parameters derived from our simulation analysis to our data (T_(10)_= 1.81, p < 0.05, cluster size = 8)^31,38^. This optimized significance map is a primary output for subsequent segmentation analysis to interpret voxel-wise information in terms of brain state anatomy.

**Fig. 4.**
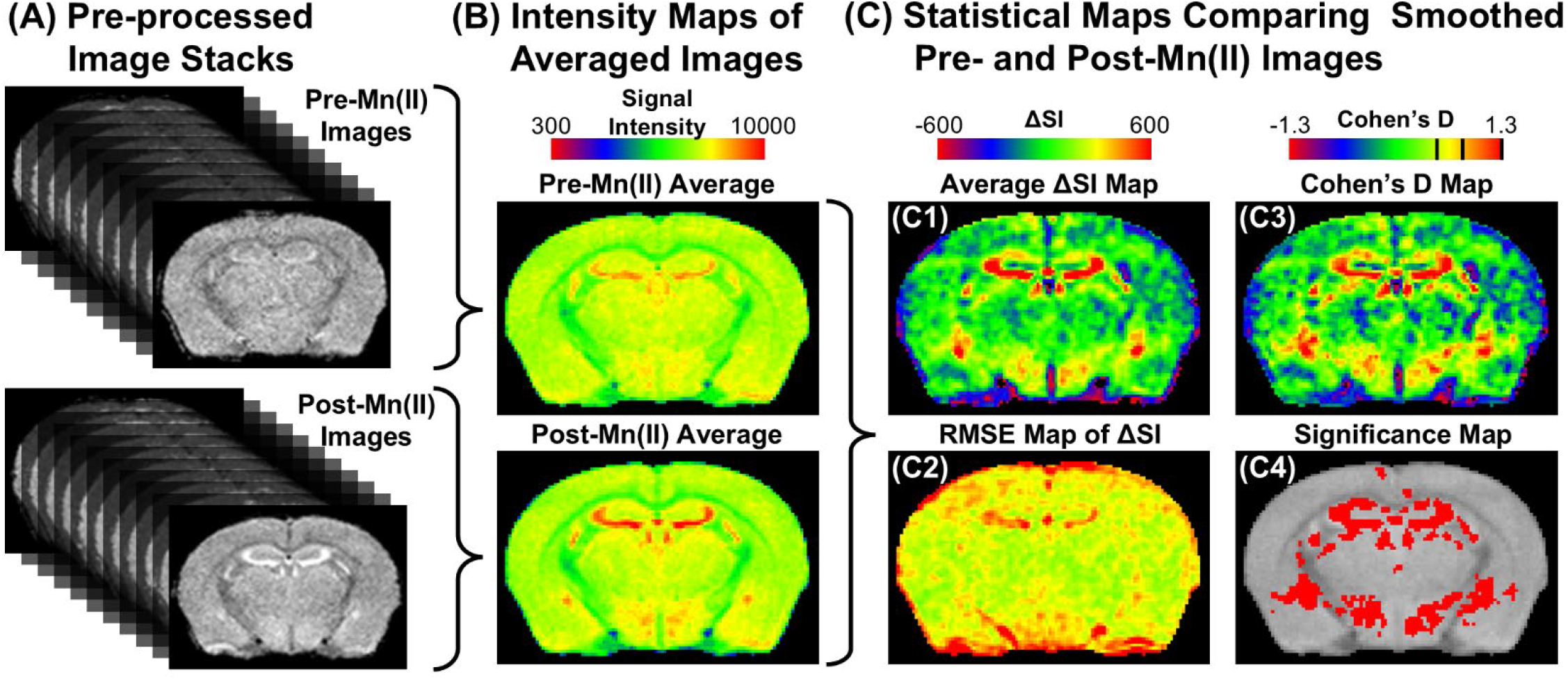
Brain-wide voxel-wise statistical mapping of Mn(II) enhancements. **(A)** Coronal slices of example data stacks of pre-Mn(II) (top) and post-Mn(II) (bottom) images after preprocessing. **(B)** Intensity maps of the average signal intensity (SI) from a single coronal slice of 11 processed pre-Mn(II) (*top*) or post-Mn(II) (*bottom*) images. FSLeyes Free Surfer color gradient^100^: Minimum and maximum set according to the average dataset noise and the histogram of averaged pre-Mn(II) image intensities. **(C)** Four different types of statistical maps comparing post-Mn(II) > pre-Mn(II) images. Signal intensity difference (ΔSI) maps of **(C1)** average voxel-wise ΔSI between post- and pre-Mn(II) images and of **(C2)** the root mean square error (RMSE) of voxel-wise ΔSI. Color gradient range approximates ± 3 St. Dev. in the averaged ΔSI image. **(C3)** Cohen’s D voxel-wise effect size map of post-Mn(II) > pre-Mn(II). Color gradient range represents Cohen’s D (d) values. From left to right, black lines in gradient indicate d = 0.5 (moderate), d = 0.8 (large), and d = 1.3 (very large). **(C4)** Statistical “significance” map of post-Mn(II) > pre-Mn(II) comparison. Red voxels are those reaching statistical significance at p < 0.05 with cluster correction for multiple comparisons; cluster size = 8, T_(11)_ = 1.81; d > 0.54. Statistical maps in (C) were masked in GIMP using a binarized dataset MDT image, with a SI threshold of 4000.

### 3.4. InVivo segmentation software for brain state characterization

To segment MEMRI data, we aligned the *InVivo* Atlas grayscale and label images to the MDT using inverse alignment procedures from different MR-focused software (SPM, FSL, and ANTs) (**Fig. 5**). We assessed alignment quality between the dataset and the *InVivo* Atlas using both visual inspection (**Fig. 5B**) and NMI (**Fig. 5C**). Each software produced strong agreement between the Atlas and our MDT (NMI > 0.8), and minimal alignment variability between individuals (COV_NMI_≈ 2.8 - 2.9%). Thus, the *InVivo* Atlas can be accurately aligned to an MR dataset using typical inverse procedures, which enables segmentation. Once aligned, the newly developed InVivoSegment software, which can be run from a GUI (**Fig. 6**), applies segment labels to MR datasets with a variety of experimental designs, and calculates measures of segmental volume and intensity.

**Fig. 5.**
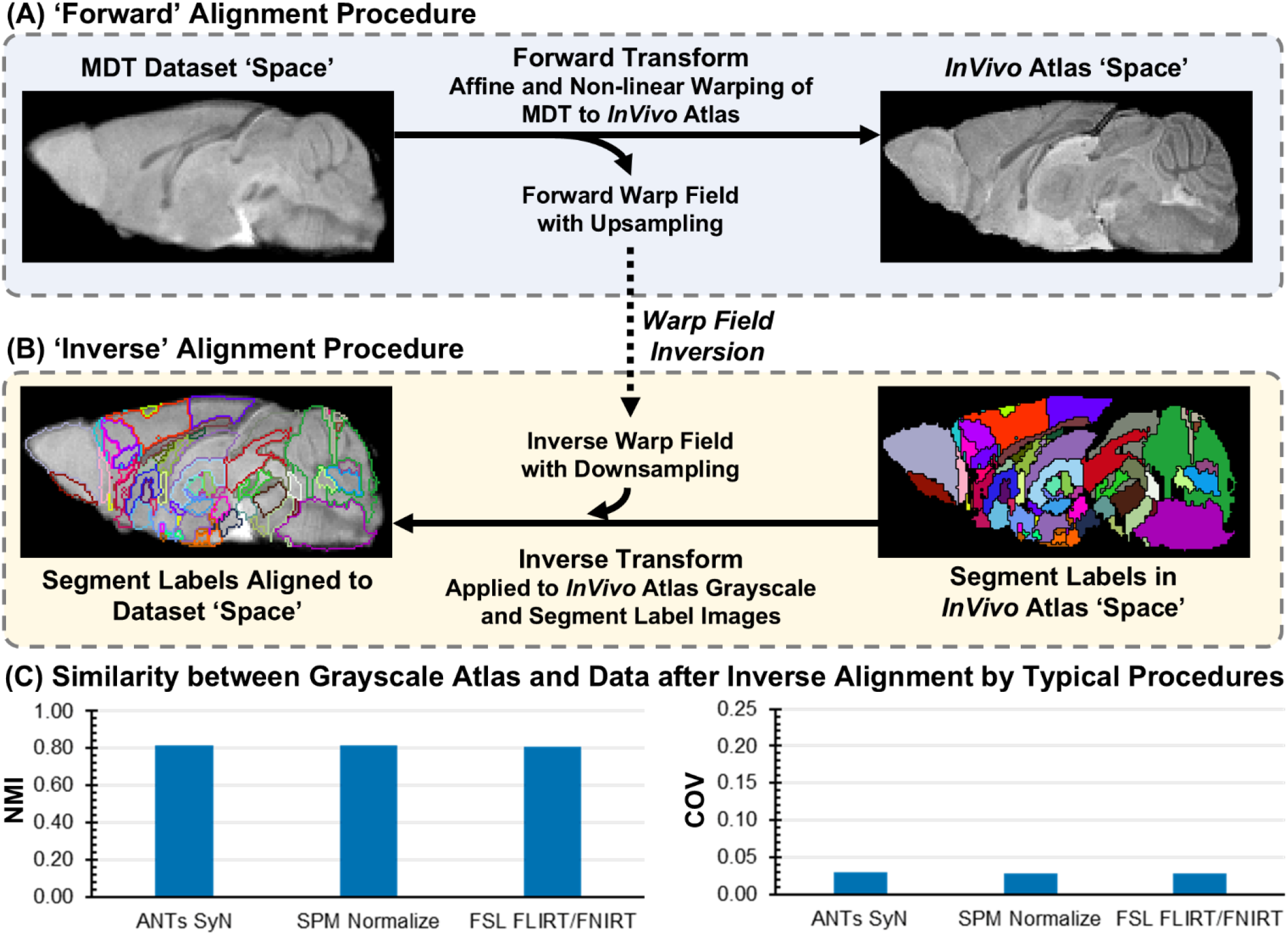
Registration of *InVivo A*tlas to a dataset template via inverse alignment. **(A) ‘**Forward’ Alignment Procedure: a lower resolution MDT image (*left*, 100 μm^3^) undergoes a multistep linear-nonlinear spatial registration to the higher resolution *InVivo* Atlas grayscale image (*right*, 80 μm) with voxel upsampling to generate a ‘forward’ warp field. Many image processing software (e.g., ANTs, FSL, SPM) provide multi-stage alignment algorithms, which first perform a 12 d.o.f. affine alignment followed by localized nonlinear warping^7,9,60^. **(B)** Inverse Alignment Procedure: Warp/deformation fields characterizing the forward transformation are inverted and then applied to the *InVivo* Atlas grayscale and label images with downsampling into the dataset space. **(C)** Bar charts of NMI for alignment quality of the *InVivo* Atlas grayscale image to our dataset using three commonly used software (ANTs, SPM, and FSL). *Left*: NMI between the *InVivo* Atlas grayscale image and the MDT. *Right:* COV of NMI between individual images after warping to the MDT and the *InVivo* Atlas grayscale image.

**Fig. 6.**
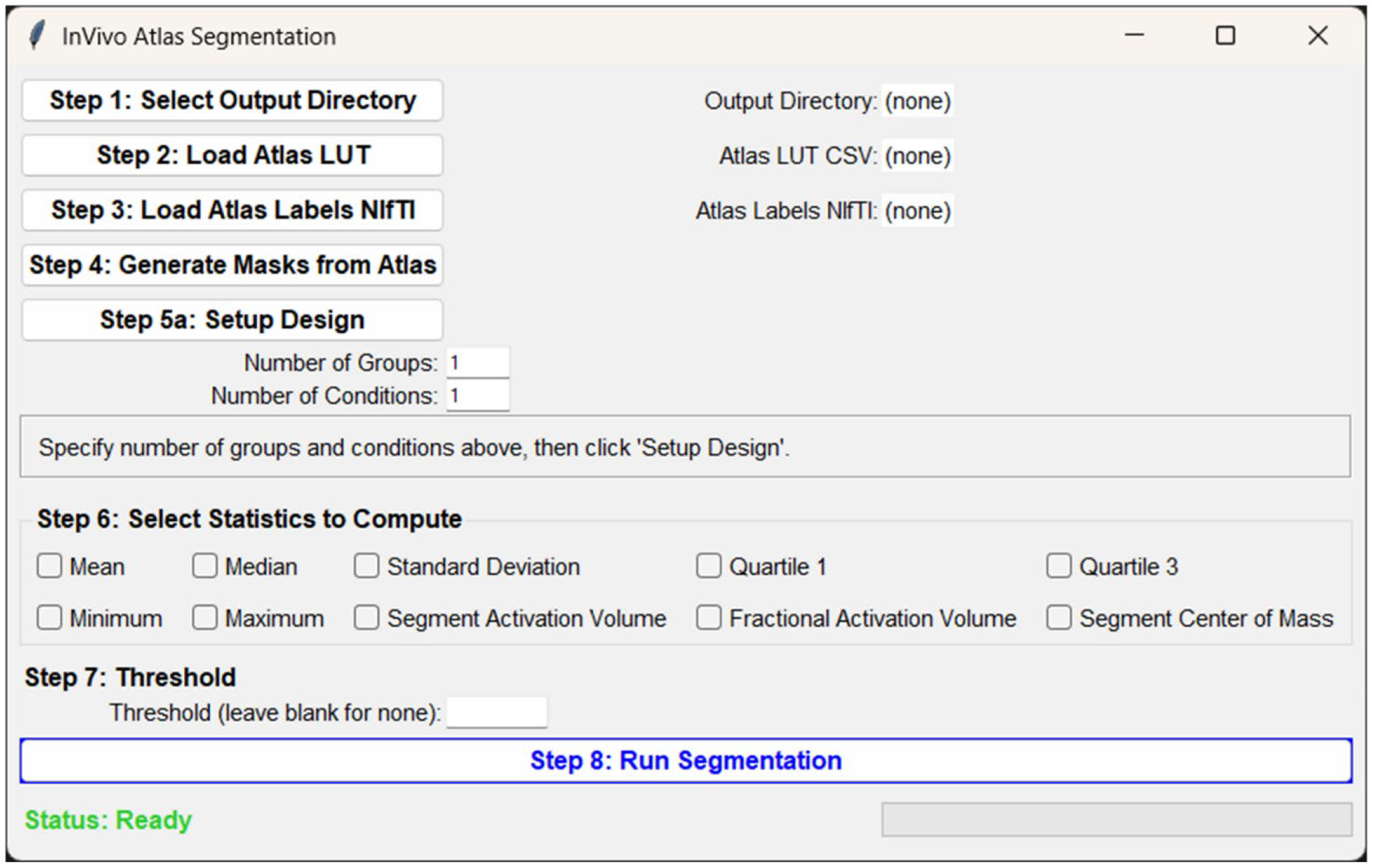
Diagram of the InVivoSegment GUI. The segmentation GUI enables application of *InVivo* Atlas labels to neuroimaging datasets (NIfTI file types) using Python. The software utilizes a reference brain atlas, both a grayscale MR image and anatomical labels in NIfTI format, and a corresponding look-up table (LUT) for organization and naming. The software allows flexible ‘Group × Condition’ experimental designs with different sample sizes and can be applied to a variety of different species or types of MR images (e.g., human or animal, T1 or T2, enhanced or not) given an appropriate Atlas/LUT is available in the correct format. Ten different measures of segmental volume and intensity are available, calculated using suprathreshold (step 7) voxels. These measures together with segment abbreviation, group, condition, subject variables populate an output CSV.

To validate the InVivoSegment software, we simulated an image with 4 signal clusters of known intensity distributions in 3 different brain segments and compared segmental measurements by InVivoSegment to those obtained from investigator calculation (**Supplemental Fig. S2**). Measures of segmental volumes and intensities (Mean, Min, Max, Q1, Median, Q3, SD, ActVol, Fractional Activation Volume [FAV], and signal-weighted centroids [x/y/z/]) were calculated at 3 different intensity thresholds. For each threshold, the InVivoSegment software reproduced investigator results (**Supplemental Table S3**).

### *3*.*5. InVivo* Atlas segmentation of real MEMRI data

As an example on real MEMRI data, we applied the InVivoSegment software to the effect-size map of Mn(II)-accumulations from Fig. 4C, which used the optimized t-value and cluster-size parameters (**Fig. 7A**). This new analysis provides more precise and deeper understanding of images we reported in ^31,38^. We further filtered segments from MEMRI data to only include those with FAV above the Beta-fitted 99^th^ percentile of FAV (11.6%) calculated from statistical maps of noise-only simulations (**Fig. 7B1**). Segment-wise FAV values from Mn(II)-accumulation exceeded this threshold in 69 of 106 segments, with 4 domains containing multiple segments with FAV exceeding 50% (**Fig. 7B1**). This demonstrates robust Mn(II)-induced signal enhancements throughout the brain. Measures of voxel intensity (effect sizes) further differentiate segmental activity patterns. For instance, Cohen’s D effect-sizes for olfactory and hippocampal segments were larger than most other segments (**Fig. 7B2**). Of the 69 segments meeting the FAV threshold, 16 displayed a slight inhomogeneity of effect size magnitudes (e.g., COV > 0.3) (**Fig. 7B3**). Moreover, a few segments exhibited noncentrally localized activation volumes, including a slight lateralization in amygdalar segments and a dorsoposterior shift in the piriform cortex (**Fig. 7B4**), suggesting potential subsegmental activations. Thus, by combining information from multiple segmental measures, a more detailed understanding of brain state can be obtained.

**Fig. 7.**
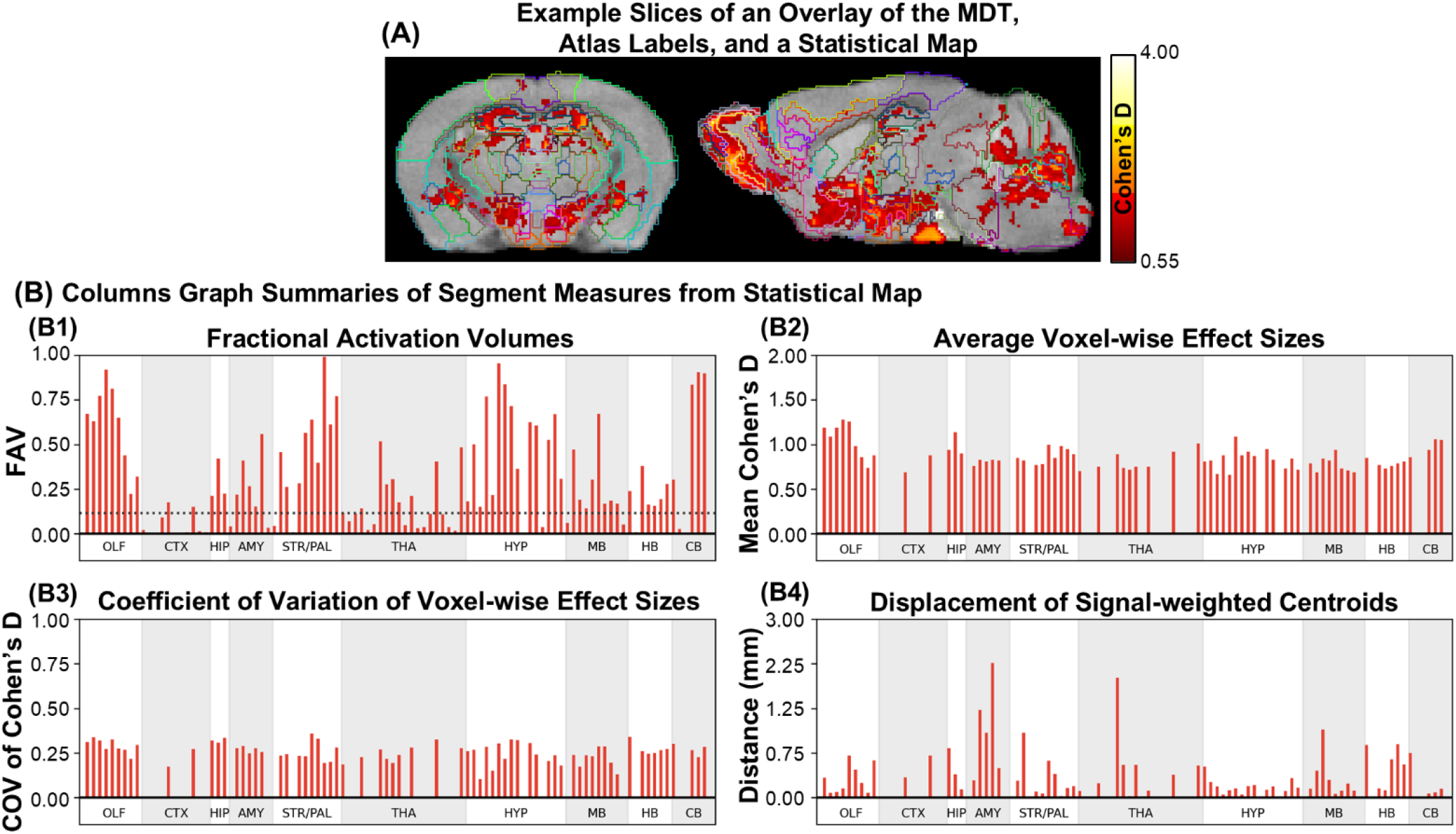
Segment-wise summaries of voxel-wise statistics to characterize brain state. **(A)** Examples of brain slices of the MDT (*left*, coronal; *right*, sagittal) with a masked Cohen’s D effect size map and *InVivo* Atlas labels overlaid. Color gradient represents voxel effect sizes ranging above statistical significance by paired t-test (n = 11, T_(10)_ ≥ 1.812, d > 0.55-4.00, p < 0.05, cluster *corr*.). **(B)** Four types of segment-wise statistical summaries calculated from *InVivo* Atlas segmentation, shown as column graphs. **(B1)** Fractional activation volumes (FAV), reflecting activation volume normalized by the size of that region. **(B2)** Average Cohen’s D effect-size of voxel-wise signals, measuring standardized magnitude of enhancement across the sample. **(B3)** COV of voxel-wise effect sizes, reflecting the degree variability or uniformity of effect magnitudes within a segment. **(B4)** Euclidean distance of the signal-weighted centroid from the volumetric centroid of that segment, reflecting subsegment localization of activation volumes. Columns represent 106 segments from 10 different anatomical domains (Olfactory, OLF; Cortex, CTX; Hippocampus, HIP; Amygdala, AMY; Striatum-Pallidum, STR/PAL; Thalamus, THA; Hypothalamus, HYP; Midbrain, MB; Hindbrain, HB; and Cerebellum, CB). For full list of segments and ordering, see **Supplemental Table S1**.

## 4. Discussion

### 4.1. Considerations for QA when preprocessing MEMRI datasets

The MEMRI processing pipeline with integrated QA proposed here provides clear checkpoints and expectations for quantitative readouts of image quality from MR acquisition to non-linear warping. Processing steps used here for mouse parallel standards developed for other species ^1-9,11^. MEMRI signal intensities reflect a combination of neural activity and connectivity as well as pharmacokinetics^15^, as compared to pure anatomical contrast. Consequently, experimental design and processing steps for MEMRI must consider sources of signal intensities to ensure that meaningful neurophysiological signals are preserved^71^. For review of experimental considerations^15,72-75^. Researchers should be cautious with bias-field corrections^76,77^, which can misinterpret ‘low-frequency’ Mn(II)-enhancement as artifact. Alignment procedures must preserve intensity concentrations when applying deformations and re-slicing^60,78^. In datasets with greater anatomical variability (e.g., genetic knockouts or neurodegenerative studies), regions with large standard deviation of deformation magnitude (SDDM) should be carefully inspected to determine whether further skull-stripping or alignment optimization is needed, or whether voxel-wise comparison is even possible. Alternative normalization choices such as phantoms^79^, background noise, other brain regions^80^or muscle^81^could be useful, but could hinder sensitivity or specificity by detection of signals from non-neural sources. Although quantitative metrics provide objective measures of image quality and processing performance, visual inspection is still a critical complement to quantitative QA^42^.

### 4.2. Utilizing simulation and balanced accuracy to optimize smoothing, effect size, and cluster size parameters

Our findings reveal three general criteria for optimizing smoothing kernel-, effect- and cluster-size parameters. First, apply the minimal 1.5x voxel smoothing kernel to meet SPM assumptions. While the Matched Filter Theorem suggests kernel sizes should be matched to the expected cluster-size of an activation^82,83^, brain-wide Mn(II) accumulations are of varying, unknown sizes, and thus a minimal smoothness is preferred. Second, use the minimal effect-size threshold to meet the FPR and FNR determined for *a priori* power analysis and sample size requirements. This selection maintains sensitivity to lower activity levels that might typically be ignored by stringent FPR correction strategies. Lastly, apply a cluster-size threshold that is close to the resolution of the data, e.g., 8 - 27 voxels for data smoothed by a 1.5x voxel smoothing kernel. This removes noise-based clusters that can occur from smoothing, while retaining spatial specificity.

A limitation of this parameter specification strategy is that it was optimized for the MR-noise and variance specific to our 11.7T MR data and our large, n = 11, sample size. While this approach could be used to determine SPM parameters in general, they must be optimized for each specific dataset. If the experimental goals were to characterize only the most specific signals from large activations, larger smoothing kernels, effect-sizes, and cluster-sizes would be reasonable selections. An alternative goal might be to optimize the balance between false discovery rate (false positives over total positives) with sensitivity. In this case, our results suggest using slightly larger cluster- and/or effect-size thresholds (d = 0.546 and cluster size = 125; or d = 0.806 and cluster size = 27). Non-parametric approaches also provide robust alternatives^61,84^. Here, we used balanced accuracy and Youden’s J from standard receiver operator characteristic (ROC) methodology, as these metrics balance sensitivity and specificity explicitly and are robust to imbalanced data^85,86^. Weighted metrics^87^ or alternative ROC approaches that use family-wise error^84,88^ may also be useful. Similar percentages of balanced accuracy may arise from different combinations of FPR and FNR. Examining the ratio of false positives removed to false negatives added by a correction strategy further informs optimization. We recommend customizing parameter optimization for individual datasets based on the underlying hypotheses being addressed and to report results at multiple levels^89^.

### 4.3. Voxel- and segment-wise statistical summaries

Our results also demonstrate three major considerations when segmenting MEMRI data: 1) large non-zero differences in aggregated statistics across multiple segments are highly likely to reflect large-scale differences in Mn(II)-enhancement and not noise; 2) segment-level FAV exceeding those produced by noise-only simulations (e.g., 11.7%) are more likely to reflect true signals; and 3) signal-based segmental measures of voxel-wise statistical maps should be interpreted jointly with FAV since small numbers suprathreshold voxels can influence their magnitudes disproportionately.

Previously, FAV highlighted both large-scale and localized dynamics in response to an acute predator threat experience^31^, and identified differences in those dynamics over time after early life adversity^38^. Deeper dives into FAV and other signal based segmentation statistics, may even inform on cell-type and circuit specific activity within brain regions^90,91^. Although not included in the current software, other statistics such as, skewness and kurtosis, or comparative metrics like Wasserstein distance and Kullback–Leibler divergence, could provide further quantitative readouts about subsegmental signal distributions^92^. Moreover, analyses that measure covariance of signal intensities between segments, including principal or independent component analysis, structural equation modeling, and graph theory, have potential to reveal details on network-level coordination of activity patterns revealed by statistical mapping^29,44,93,94^.

Limitations of this segment-wise analysis of voxel-wise intensities include differing volumes of segmented regions, activation volumes overlapping multiple segments, and multiple subsegmental activations, among others. While volumetric biased activity can be mitigated somewhat using FAV, activations in larger segments and multiple subsegmental activations may still go unidentified. An alternative approach to our anatomical segmentation would be to segment the brain by arbitrarily specified uniform voxel clusters, e.g.., image partitioning or tokenization, and/or by unsupervised segmentation of functional signals^95,96^. Small amounts of activity (volume or intensity) may have disproportionately large effects on brain state, and may not be easily assigned to a behavioral outcome^97,98^, whereas large amounts of activity may be lost in homeostatic noise (motor output, metabolism/feeding, etc.).

## 5. Conclusion

We propose an integrated preprocessing pipeline that can lead to standardization of mouse brain MR analysis, with data-driven quality assurance, simulation-based validations, and optimization strategies for selection of key parameters for voxel-wise statistical mapping that balance sensitivity with specificity. We provide methods for creating simulated image matrices for parameter optimization and a GUI for automated segmentation of mouse MR data based on our high-resolution and Mn(II)-enhanced *InVivo* Atlas, and multiple examples for individual validation. This pipeline and associated software tools were developed with the intent to provide clear steps for quality assurance and for verification and validation of brain-wide MR image analysis; a scalable segmentation framework for mouse MR data; and to facilitate cross-study comparison. Please let us know if you find this new pipeline useful!

## Acknowledgements

We gratefully acknowledge support from the Biological Imaging Facility/Center and the Beckman Institute at Caltech where imaging was originally performed. We thank the staff, faculty and trainees at UNM, USC and Caltech for research and administrative support. This work was funded by the National Institutes of Health R01MH096093 (Bearer), P30AG08404 (Rosenberg/Bearer), P20AG068077 (Rosenberg/Bearer), F99NS139535 (Uselman); Zilkha Neurogenetic Institute (Jacobs); Beckman Institute at Caltech (Bearer and Jacobs); and Harvey Family Endowment at UNM (Bearer).

## Data and Code Availability

MEMRI data analyzed in this manuscript were published previously with OpenNeuro and available at https://openneuro.org/datasets/ds006746/versions/1.0.0. Code used in this manuscript for MR data processing and analysis is available at https://github.com/bearerlab/memri-processing-QA. The InVivoSegment software package and associated *InVivo* Atlas files are available at https://github.com/bearerlab/InVivoSegment.

