## Supplemental Information - Methods and Figures for "Quality Assurance Strategies for Brain State Characterization by MEMRI"

**This file contains:**

1. **Supplemental Methods: Formulation of algorithm for simulated true positive data.**
2. **Supplemental Figures S1 - S2**
3. **Supplemental Tables S1 - S3**

**1. Supplemental Methods.**

**1.1. Formulation of algorithms for simulating false- and true-positive images.**

To simulate true positive signals with targeted sample-wise effect sizes of $\boldsymbol{d}$ = 0.546, 0.801, and 0.950, we simulated intensity differences based on a Gaussian distribution parameterized by the targeted mean difference $\boldsymbol{\mu_\Delta}$ and brain-wide average voxel-wise signal variance $\boldsymbol{v} = (\boldsymbol{\sigma}^2 + \boldsymbol{\varepsilon}^2)$. Note that $\boldsymbol{v}$ was determined from the variance image produced by SPM paired t-test on unsmoothed data; $\boldsymbol{\sigma}$ = sd of inter-individual pairwise voxel-wise intensity differences between pre- and post-Mn(II) images, which was estimated as $\boldsymbol{\sigma} = \sqrt{\boldsymbol{v}^2 - 2\boldsymbol{\varepsilon}^2}$); and that $\boldsymbol{\varepsilon}$ = average voxel-wise noise derived from SNR measurements. We simulated $\boldsymbol{N}$ = 10,000 samples of size $\boldsymbol{n}$ = 11, calculated the effect size for each sample, and selected the realization that minimized the difference between simulated and targeted effect-size. True positive intensity differences from the optimal realization were extracted and embedded into the 3D lattice of voxel clusters. This process is formulated below in [**Eq. 1**] - [**Eq. 5**].

**Step 1 - Parameterize and derive the targeted mean difference $\boldsymbol{\mu_\Delta}$:**

$$\boldsymbol{\mu_\Delta} = \boldsymbol{d} \cdot \boldsymbol{\sigma} \tag{1}$$

**Step 2 - Simulate N paired difference samples:**

For each realization $\boldsymbol{i} = \mathbf{1}, \dots, \boldsymbol{N}$, with sample size $\boldsymbol{n} = \mathbf{11}$.

$$\mathrm{x}^{(i)} = \{\boldsymbol{x}_1^{(i)}, \dots, \boldsymbol{x}_n^{(i)}\} \sim \mathcal{N}(\boldsymbol{\mu_\Delta}, \boldsymbol{v}) \tag{2}$$

**Step 3 - Compute the sample Cohen's d for each realization:**

$$\widehat{\boldsymbol{d}}^{(i)} = \frac{\bar{\boldsymbol{x}}^{(i)}}{\boldsymbol{s}^{(i)}} \tag{3}$$

where $\bar{\boldsymbol{x}}^{(i)}$ and $\boldsymbol{s}^{(i)}$ are the sample mean and standard deviation of $\mathrm{x}^{(i)}$.

**Step 4 - Select the optimal realization $\boldsymbol{i}^*$ and extract the simulated intensity values $\mathrm{x}^{(i^*)} = \{\boldsymbol{x}_1^{(i^*)}, \dots, \boldsymbol{x}_n^{(i^*)}\}$:**

$$\boldsymbol{i}^* = \boldsymbol{argmin}_{\,i \in \{1,\dots,N\}} \left|\widehat{\boldsymbol{d}}^{(i)} - \boldsymbol{d}\right| \tag{4}$$

**Step 5 - Assemble pairwise signals $\mathrm{s}_1$ and $\mathrm{s}_2$ for each paired set of images $\boldsymbol{j}$ into 3D lattice of voxel clusters:**

$$\boldsymbol{s}_{2,\mathrm{j}}^{(i^*)} = \boldsymbol{s}_{1,j} + \boldsymbol{x}_{\mathrm{j}}^{(i^*)}, \boldsymbol{j} = \mathbf{1}, \dots, \boldsymbol{n} \tag{5}$$

**2. Supplemental Figures.**

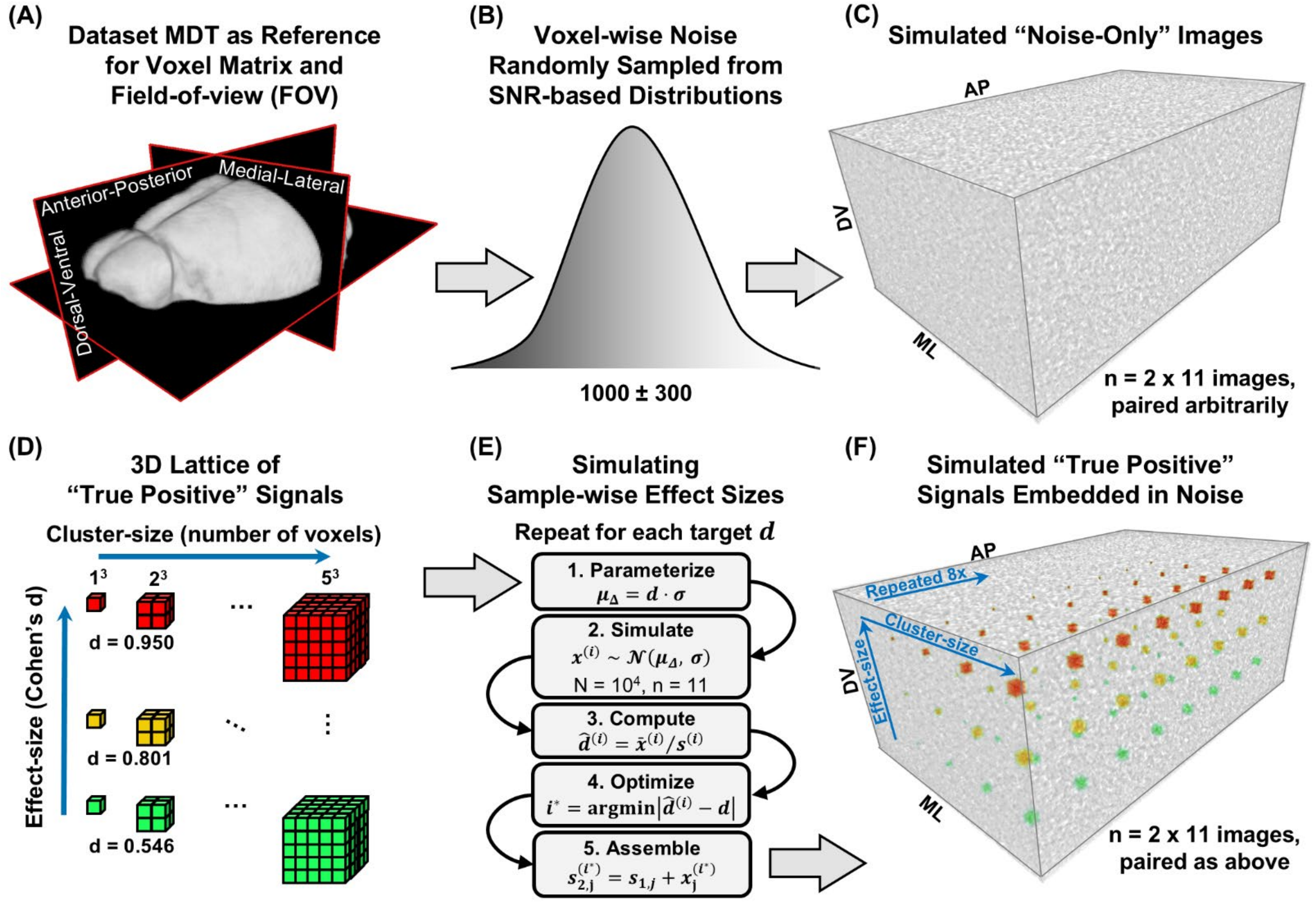


**Fig. S1. Process for generating simulated "noise-only" and "true-positive" images. (A)** A 3D rendering of the dataset MDT intersected by orthogonal slices from each anatomical plane (black rectangles). The MDT's voxel matrix and field-of-view (FOV) were used as a template in which to embed simulated noise and/or positive signals. **(B)** A representation of the grayscale intensity distribution from which MR noise was sampled independently for each voxel in the 22 images of the simulated dataset. The noise signal distribution was parameterized by a Gaussian with average and standard deviation of 1000 ± 300. A total of twenty-two images were generated, corresponding to 11 arbitrary pairs: {1, 11}, {2, 13}, …, {11, 22}. **(C)** An example noise-only image rendered in 3D using the same orientation and perspective as the MDT in (A). AP = Anterior-Posterior, ML = Medial-Lateral, DV = Dorsal-Ventral; MDT matrix/FOV boundary (dark gray outlines). Grayscale MR noise is shown as grayscale signals. **(D)** A 3D lattice of cubic voxel clusters corresponding to 3 different effect-sizes (*bottom to top;* d = 0.546, green; d = 0.801, yellow; d = 0.950, red) and 5 different cluster-sizes (*left to right*, $1^3$ - $5^3$ voxels). Each cluster is separated by empty voxels in each direction to mitigate signal overlaps from Gaussian smoothing prior to statistical mapping. The effect-by-cluster-size lattice was repeated 8 times in the third dimension (*into the page*), to generate the full 3D lattice of cubic voxel clusters. **(E)** Samples of paired true positive signals were simulated to match the expected sample-wise effect-sizes for each Cohen's d level in the 3D lattice. This processes included 5 steps: 1) Parameterize the mean difference distribution from which to sample signal differences using the targeted Cohen's d and inter-individual SD of voxel-wise signal intensity $\boldsymbol{\sigma}$; 2) Perform N = 10,000 samples of size n = 11 from this distribution; 3) Compute the effect-sizes ($\hat{\boldsymbol{d}}^{(i)}$) for each of the 10,000 samples; 4) Identify the sample ($\boldsymbol{i}^*$) that minimized the difference between the sample effect-size and targeted effect-size; 5) Compute signal intensities for each of the 11 paired images based on the optimized sample, assemble signals into the corresponding voxel clusters of the 3D lattice and embed signals into to noise-only image pairs. **(F)** A 3D rendering of same noise-only image in (C) with the lattice of true positive sample-wise effect-sizes embedded. Blue arrows indicate orientation of the 3D lattice in relation to (D). Effect-size varied along the DV axis and cluster-size along ML axis. This pattern was repeated 8 times along the AP axis. The sample signal intensity differences across all 11 image pairs yielded an effect-size map that reflects the lattice of true positives shown here.

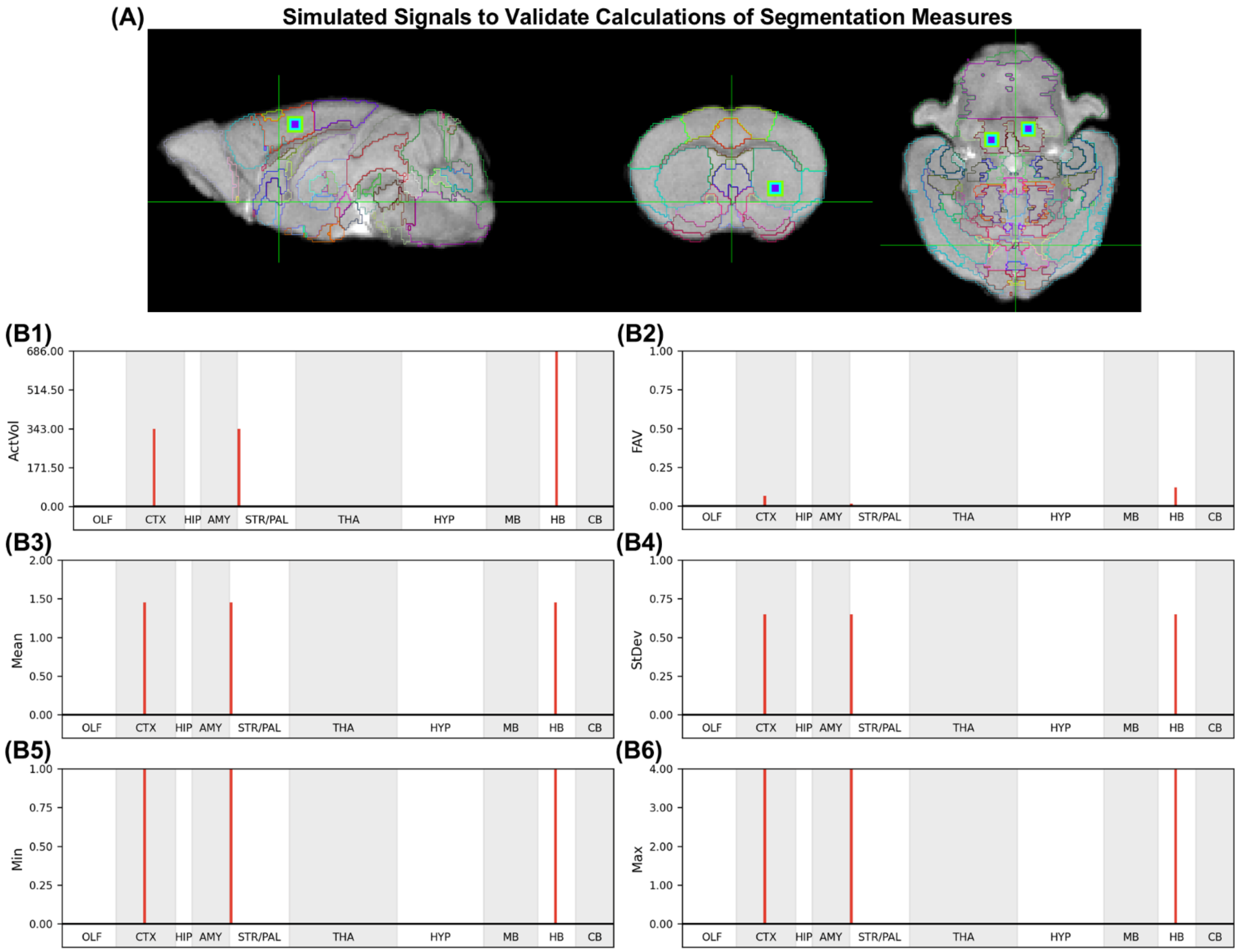


**Fig. S2. Validation for semi-automated segmentation and statistical calculations. (A)** Sagittal, coronal, and axial slices in a 3D image showing the location of 4 different 7x7x7 cubes of simulated signal for validating. One cube was placed in the ACA, one in CP,  and two in PRN. FSLeyes Free Surfer color gradient: min = 1, max = 4. The outermost shell has an intensity of 1, the second layer has an intensity of 2, the third shell has an intensity of 3, and the innermost voxel has an intensity of 4. **(B)** Column graphs of brain wide segmentation statistics calculated after segmenting the validation images based on voxel intensities > 0. **(B1)**  Log 10 number of segment voxels (SegVol, log10), demonstrate that all regions were segmented. **(B2-B6)** Segmentation statistics (FAV, mean, standard deviation, minimum and maximum) demonstrate successful detection of all three regions in which signals were simulated. For the full list of statistics calculated at different signal thresholds and how they match our secondary calculations performed in Excel, please see **Table S3** and **Supplemental Data**.

### 3. Supplemental Tables

**Table S1. Key to *InVivo* Atlas abbreviations and ordering for anatomical domains and segments.**

| Domain | Abbreviation | Segment Name |
|---|---|---|
| **Olfactory System: OLF** | | |
| OLF | MOBgl | Main olfactory bulb glomerular layer |
| OLF | MOBgr | Main olfactory bulb granule layer |
| OLF | MOBipl | Main olfactory bulb inner plexiform layer |
| OLF | MOBmi | Main olfactory bulb mitral layer |
| OLF | MOBopl | Main olfactory bulb outer plexiform layer |
| OLF | AOB | Accessory olfactory bulb |
| OLF | AON | Anterior olfactory nucleus |
| OLF | EPd | Endopiriform nucleus dorsal part |
| OLF | TT | Taenia tecta dorsal part |
| OLF | SEZ | Subependymal zone |
| **Cerebral Cortex: CTX** | | |
| CTX | CTX | Cerebral cortex (unsegmented) |
| CTX | ORB | Orbital frontal cortex |
| CTX | PL | Prelimbic cortex |
| CTX | ILA | Infralimbic cortex |
| CTX | DP | Dorsal peduncular cortex |
| CTX | ACA | Anterior cingulate cortex |
| CTX | MO | Primary motor cortex |
| CTX | SS | Somatosensory cortex |
| CTX | PIR | Piriform cortex |
| CTX | RSP | Retrosplenial cortex |
| CTX | PTL | Posterior parietal association area |
| **Hippocampus: HIP** | | |
| HIP | HPF | Hippocampal formation |
| HIP | DG | Dentate gyrus |
| HIP | CA1-CA3 | Field CA1, CA2, CA3 pyramidal layers |
| **Amygdala: AMY** | | |
| AMY | AAA | Anterior amygdala area |
| AMY | MEA | Medial amygdalar nucleus |
| AMY | CEA | Central amygdalar nucleus |
| AMY | LA | Lateral amygdalar nucleus |
| AMY | BLA | Basolateral amygdalar nucleus |
| AMY | PA | Posterior amygdalar nucleus |
| AMY | COA | Cortical amygdalar area |
| **Striatum and Pallidum: STR/PAL** | | |
| STR/PAL | CP | Caudoputamen |
| STR/PAL | ACB | Nucleus accumbens |
| STR/PAL | FS | Fundus of striatum |
| STR/PAL | OT | Olfactory tubercle |
| STR/PAL | LSc | Lateral septal nucleus caudal part |
| STR/PAL | LSr | Lateral septal nucleus rostral part |
| STR/PAL | GPe | Globus pallidus external |
| STR/PAL | SI | Substantia innominata |
| STR/PAL | MS | Medial septal nucleus |
| STR/PAL | NDB | Diagonal band nucleus |
| STR/PAL | BST | Bed nuclei of the stria terminalis |
| **Thalamus: THA** | | |
| THA | VAL | Ventral anterior lateral complex of the thalamus |
| THA | VM | Ventral medial nucleus of the thalamus |

| | | |
|---|---|---|
| THA | VPM | Ventral posteromedial nucleus of the thalamus |
| THA | VPL | Ventral posterolateral nucleus of the thalamus |
| THA | SPA | Subparafascicular area |
| THA | MG | Medial geniculate complex |
| THA | LGd | Dorsal part of the lateral geniculate complex |
| THA | LP | Lateral posterior nucleus of the thalamus |
| THA | AV | Anteroventral nucleus of thalamus |
| THA | IAM | Anteromedial nucleus of thalamus |
| THA | IMD | Intermediodorsal nucleus of the thalamus |
| THA | PVT | Paraventricular nucleus of the thalamus |
| THA | MD | Mediodorsal nucleus of thalamus |
| THA | PT | Parataenial nucleus |
| THA | RE | Nucleus of reunions |
| THA | RT | Reticular nucleus of the thalamus |
| THA | CM | Central medial nucleus of the thalamus |
| THA | PO | Posterior complex of the thalamus |
| THA | PF | Parafascicular nucleus |
| THA | MH | Medial habenula |
| **Hypothalamus: HYP** | | |
| HYP | HY | Hypothalamus (unsegmented) |
| HYP | MPO | Preoptic nuclei |
| HYP | AVPV | Anteroventral periventricular nucleus |
| HYP | PVH | Paraventricular hypothalamic nucleus |
| HYP | SCH | Suprachiasmatic nucleus |
| HYP | AHN | Anterior hypothalamic nucleus |
| HYP | SBPV | Subparaventricular zone |
| HYP | LHA | Lateral hypothalamic area |
| HYP | ZI | Zona incerta |
| HYP | ARH | Arcuate hypothalamic nucleus |
| HYP | VMH | Ventromedial hypothalamic nucleus |
| HYP | DMH | Dorsomedial hypothalamic nucleus |
| HYP | PVp | Periventricular hypothalamic nucleus (posterior part) |
| HYP | PH | Posterior hypothalamic nucleus |
| HYP | STN | Subthalamic nucleus |
| HYP | MBO | Mammillary nuclei of the hypothalamus |
| **Midbrain: MB** | | |
| MB | MB | Midbrain (unsegmented) |
| MB | VTA | Ventral tegmental area |
| MB | RN | Red nucleus |
| MB | SNc | Substantia nigra compact part |
| MB | SNr | Substantia nigra reticular |
| MB | IPN | Interpeduncular nucleus |
| MB | CLI | Central linear raphe |
| MB | CS | Superior central raphe |
| MB | DR | Dorsal nucleus raphe |
| MB | PAG | Periaqueductal gray |
| **Hindbrain: HB** | | |
| HB | P | Pons |
| HB | PG | Pontine gray |
| HB | PCG | Pontine central gray |
| HB | PRN | Pontine reticular nucleus |
| HB | PB | Parabrachial nucleus |
| HB | LC | Locus coeruleus |
| HB | MY | Medulla oblongata |
| **Cerebellum: CB** | | |

| | | |
|---|---|---|
| CB | CB | Cerebellum (unsegmented) |
| CB | DEC | Declive VI |
| CB | FOTU | Folium-tuber vermis VII |
| CB | PYR | Pyramus VIII |
| CB | NOD | Nodulus X |
| CB | UVU | Uvula IX |
| CB | SIM | Simple lobule |

**Table S2. Cluster size thresholds and effect size filtering for balancing sensitivity and specificity of statistical mapping.**

Shown are results from the false positive analysis based on voxel-wise paired *t*-tests ($p < 0.05$, $T_{(10)} \geq 1.81$) performed on noise-only simulated data after smoothing with a 1.5x voxel Gaussian kernel, corresponding to the analyses shown in **Fig 4B-D**. Power analyses using regional measurements from pilot MEMRI data indicated that a sample size of 10 -12 animals is required to achieve 80–90% power. Therefore, significant results at $p < 0.05$ represent FPR = 0.05 and FNR = 0.2 for our sample size. Bold italicized entries indicate the effect and cluster-size combination used for MEMRI analyses, which was selected to balance removal of false positives with sensitivity to small signal clusters of moderate effect size (e.g., $d > 0.546$ in ≥ 8 contiguous voxels).

***Smoothing Kernel = 1.5x voxels; SPM Paired T-test, p < 0.05***

| Cohen's D Effect Size | Cluster Size | Number of Significant Voxels | False Positive Rate | False Negative Rate | Balanced Accuracy | Yuden's J Statistic |
|---:|---:|---:|---:|---:|---:|---:|
| 0.546 | 1 | 105110 | 5.0% | 17.7% | 88.7% | 77.3% |
| ***0.546*** | ***8*** | ***71702*** | ***3.3%*** | ***18.0%*** | ***89.4%*** | ***78.7%*** |
| 0.546 | 27 | 32759 | 1.4% | 20.0% | 89.3% | 78.6% |
| 0.546 | 64 | 13435 | 0.5% | 26.4% | 86.6% | 73.1% |
| 0.546 | 125 | 7294 | 0.2% | 38.7% | 80.5% | 61.1% |
| 0.806 | 1 | 25206 | 1.1% | 57.6% | 70.6% | 41.3% |
| 0.806 | 8 | 6481 | 0.2% | 58.6% | 70.6% | 41.2% |
| 0.806 | 27 | 2685 | 0.0% | 63.2% | 68.4% | 36.8% |
| 0.806 | 64 | 1998 | 0.0% | 71.7% | 64.1% | 28.2% |
| 0.806 | 125 | 955 | 0.0% | 85.6% | 57.2% | 14.4% |
| 0.950 | 1 | 10643 | 0.5% | 87.0% | 56.3% | 12.5% |
| 0.950 | 8 | 1368 | 0.0% | 88.0% | 56.0% | 12.0% |
| 0.950 | 27 | 657 | 0.0% | 90.6% | 54.7% | 9.4% |
| 0.950 | 64 | 207 | 0.0% | 96.9% | 51.5% | 3.1% |
| 0.950 | 125 | 0 | 0.0% | 100.0% | 50.0% | 0.0% |

**Table S3. Segmentation statistics for simulated validation data.**

Shown are results from semi-automated segmentation of the three segments (ACA, CP, and PRN) in which 7x7x7 voxel regions of simulated signal were placed at three different signal thresholds ( >0, >1, or >2). Statistics calculated via the segmentation GUI match those calculated via manually via Excel (see **Supplemental Data**). 'Active' voxels (also ActVol from InVivoSegment) indicate the number of voxels exceeding the threshold. sCOGx/y/z – signal-weighted centroids based on voxel location. X = medial lateral; Y = anterior-posterior; Z = dorsal-ventral. FAV, Mean, StDev, Min and Max values for a threshold > 0 correspond to values shown in column graphs in **Fig. S2**.

| Segment | Threshold | Mean | Median | StDev | Q1 | Q3 | Min | Max | Active Voxels | FAV | sCoGx | sCoGy | sCoGz |
|---|---|---|---|---|---|---|---|---|---|---|---|---|---|
| ACA | 0.0 | 1.45 | 1.0 | 0.65 | 1.0 | 2.0 | 1.0 | 4.0 | 343 | 0.067 | 62.0 | 71.0 | 60.0 |
| ACA | 1.0 | 2.22 | 2.0 | 0.44 | 2.0 | 2.0 | 2.0 | 4.0 | 125 | 0.024 | 62.0 | 71.0 | 60.0 |
| ACA | 2.0 | 3.04 | 3.0 | 0.19 | 3.0 | 3.0 | 3.0 | 4.0 | 27 | 0.005 | 62.0 | 71.0 | 60.0 |
| CP | 0.0 | 1.45 | 1.0 | 0.65 | 1.0 | 2.0 | 1.0 | 4.0 | 343 | 0.015 | 42.0 | 63.0 | 32.0 |
| CP | 1.0 | 2.22 | 2.0 | 0.44 | 2.0 | 2.0 | 2.0 | 4.0 | 125 | 0.006 | 42.0 | 63.0 | 32.0 |
| CP | 2.0 | 3.04 | 3.0 | 0.19 | 3.0 | 3.0 | 3.0 | 4.0 | 27 | 0.001 | 42.0 | 63.0 | 32.0 |
| PRN | 0.0 | 1.45 | 1.0 | 0.65 | 1.0 | 2.0 | 1.0 | 4.0 | 686 | 0.119 | 63.5 | 115.5 | 26.0 |
| PRN | 1.0 | 2.22 | 2.0 | 0.44 | 2.0 | 2.0 | 2.0 | 4.0 | 250 | 0.043 | 63.5 | 115.5 | 26.0 |
| PRN | 2.0 | 3.04 | 3.0 | 0.19 | 3.0 | 3.0 | 3.0 | 4.0 | 54 | 0.009 | 63.5 | 115.5 | 26.0 |
